# Species’ traits and exposure as a future lens for quantifying seabird bycatch vulnerability in global fisheries

**DOI:** 10.1101/2021.05.24.445472

**Authors:** Cerren Richards, Robert S. C. Cooke, Diana E. Bowler, Kristina Boerder, Amanda E. Bates

## Abstract

Fisheries bycatch, the incidental mortality of non-target species, is a global threat to seabirds and a major driver of their declines worldwide. Identifying the most vulnerable species is core to developing sustainable fisheries management strategies that aim to improve conservation outcomes. To advance this goal, we present a preliminary vulnerability framework that integrates dimensions of species’ exposure, sensitivity, and adaptive capacity to fisheries bycatch to classify species into five vulnerability classes. The framework combines species’ traits and distribution ranges for 341 seabirds, along with a spatially resolved fishing effort dataset. Overall, we find most species have high vulnerability scores for the sensitivity and adaptive capacity dimensions. By contrast, exposure is more variable across species, and thus the median scores calculated within seabird families is low. We further find 46 species have high exposure to fishing activities, but are not identified as vulnerable to bycatch, whilst 133 species have lower exposure, but are vulnerable to bycatch. Thus, the framework has been valuable for revealing patterns between and within the vulnerability dimensions. Still, further methodological development, additional traits, and greater availability of threat data are required to advance the framework and provide a new lens for quantifying seabird bycatch vulnerability that complements existing efforts, such as the International Union for Conservation of Nature (IUCN) Red List.

## Introduction

As of 2018, the global fishing fleet is estimated at 4.56 million fishing vessels of various sizes (FAO 2020). Fisheries bycatch, the incidental mortality of non-target species, is a serious threat to seabirds, driving seabird population declines worldwide (Dias et al. 2019). Thus, key goals for successful fisheries management and conservation are to identify vulnerable non-target species and develop bycatch mitigation strategies. Yet, these goals pose global challenges because seabirds are wide ranging and encounter fishing activities in various national and international waters at different stages of their life history (Komoroske and Lewison 2015). Better understanding of the factors affecting vulnerability of species to bycatch is an essential step towards predicting which species are most at risk and working to mitigate bycatch threats.

While seabird bycatch is widespread, a global quantification of seabird vulnerability to fisheries bycatch in multiple gear types (e.g. longline, trawl and purse seine) is lacking because bycatch data are scarce (Anderson et al. 2011, Hedd et al. 2016, Suazo et al. 2017, Zhou et al. 2019). There is very low observer coverage aboard fishing vessels, and existing data has poor species discrimination and only coarse quantification (Bartle 1991, Weimerskirch et al. 2000, Sullivan et al. 2006, Anderson et al. 2011, Hedd et al. 2016, Suazo et al. 2017). Thus, bycatch mortality of high-risk species may be undetected by on board vessels by fishers and observers, and therefore under- or unreported to databases that collate species’ threat data such as the International Union for Conservation of Nature (IUCN) Red List (iucnredlist.org). Coupling traits with fisheries exposure information could offer a complementary lens to existing methods and provide insights into different dimensions of seabird bycatch vulnerability.

Trait-based approaches have emerged as being important for advancing conservation efforts (Miatta et al. 2021), where traits represent fundamental biological attributes of organisms measured at the individual level (Violle et al. 2007, Gallagher et al. 2020). Furthermore, selecting ecologically meaningful and interpretable traits can relate to species’ vulnerabilities to threats (Zhou et al. 2019, Richards et al. 2021). As an exceptionally well-studied group, detailed information is available on the life history, behavioural and ecological traits of seabirds for predictive trait-based analyses (Tavares et al. 2019, Richards et al. 2021). Thus, integrating freely available global threat datasets with species traits in a vulnerability framework may be a valuable tool to identify the seabird species most vulnerable to gear-specific bycatch.

A species’ vulnerability to bycatch is determined by both extrinsic (threats) and intrinsic (traits) factors. Specifically, such factors include the interplay between a species’ exposure, sensitivity, and capacity to adapt in response to bycatch (Foden et al. 2013, Potter et al. 2017, Butt and Gallagher 2018). Firstly, exposure encompasses the extent to which species’ ranges overlap with fishing activities and the magnitude of activities experienced. For example, wide-ranging pelagic foragers, such as albatrosses, overlap with a variety of fishing gears and fleets throughout their lives (Clay et al. 2019). Secondly, sensitivity traits represent a species’ likelihood of bycatch mortality when it interacts with fisheries. For example, large seabirds have a greater risk of bycatch mortality than smaller seabirds (Zhou et al. 2019). Finally, adaptive capacity traits describe the ability for populations to adapt and recover from bycatch mortalities. For example, bycatch will have a greater impact on seabirds with slow reproductive rates, such as albatross and auks, which lay a single egg per season and reach sexual maturity after five to ten years.

Coupling a dataset of traits with seabird global range maps and a spatially resolved gear-specific fishing dataset could provide a new lens for quantifying seabird bycatch vulnerability that would complement existing efforts, such as the IUCN Red List. Here we (1) develop a framework for quantifying seabird bycatch vulnerability to multiple gear types; (2) analyse the emerging patterns of seabird bycatch vulnerability based on available data and traits; and (3) discuss future directions and visions for the vulnerability framework.

## Building a vulnerability framework

Here we modify a framework that has previously been applied to a diversity of species from birds and trees to amphibians and corals (Foden et al. 2013, Potter et al. 2017), with the goal to identify the seabird species most vulnerable to gear-specific bycatch (Fig. 1). Our intention is for the vulnerability framework to be built upon and improved as more trait and threat data become available in the future.

**Figure 1.**
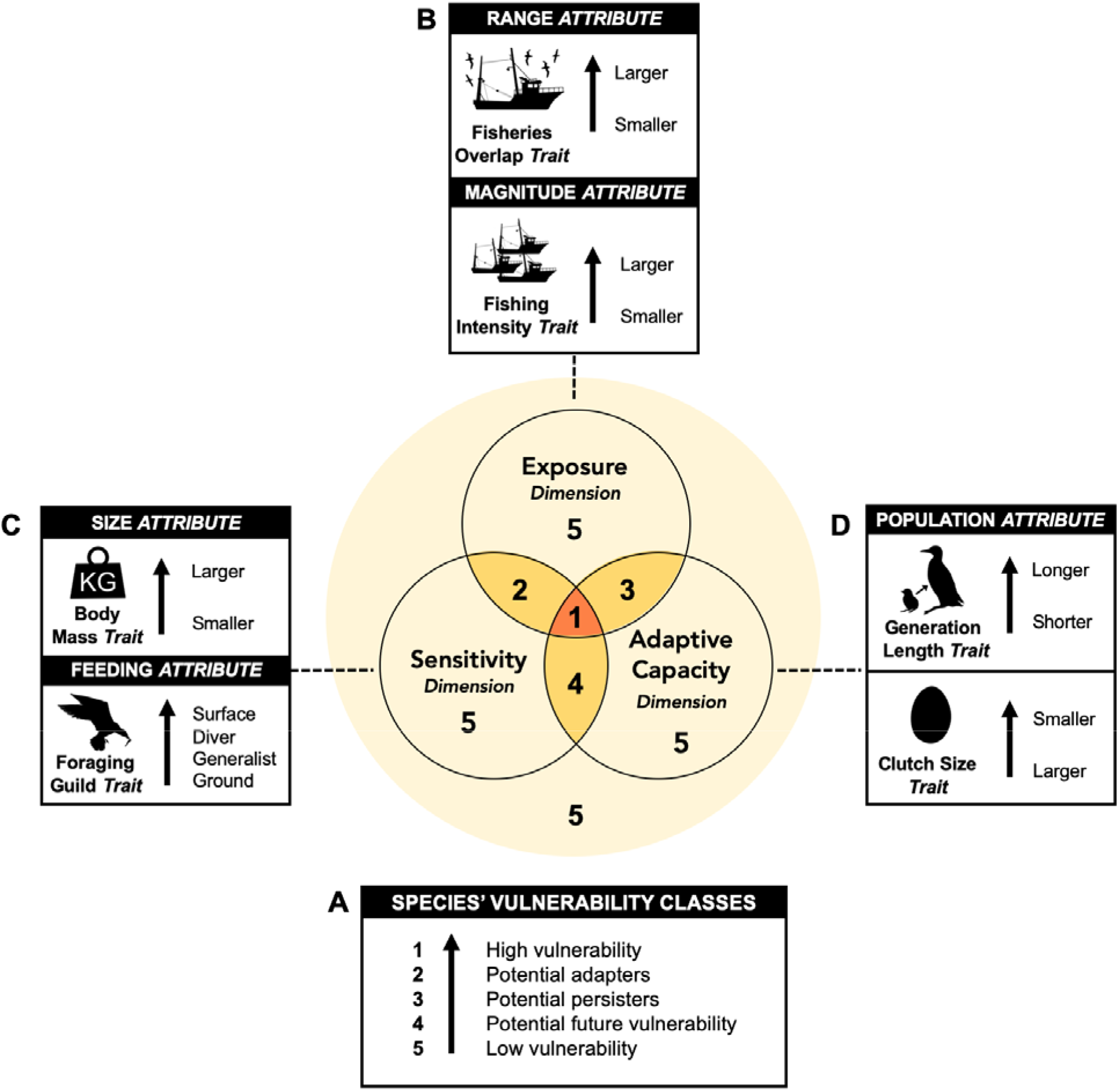
Framework to quantify species’ vulnerability to bycatch. The combination of three dimensions: exposure, sensitivity and adaptive capacity, characterise five distinct species’ vulnerability classes (Box A). Six traits associated with five overarching vulnerability attributes (Boxes B-D: Size, Feeding, Range, Magnitude, and Population) are used to quantify each vulnerability class. Black arrows indicate the direction of increased vulnerability. Modified from Foden et al. (2013) and Potter, Crane & Hargrove (2017).

The trait-based framework integrates three dimensions of bycatch vulnerability based on exposure, sensitivity, and adaptive capacity. Each dimension encompasses a set of vulnerability attributes (Size, Feeding, Range, Magnitude, Population) that in turn are represented by species’ traits (Fig. 1). The framework can be used to classify species into five vulnerability classes: high vulnerability, potential adapters, potential persisters, potential future vulnerability, and low vulnerability. Each has implications for conservation prioritisation and strategic planning (Foden et al. 2013).

### Assessing sensitivity and adaptive capacity to bycatch

We selected body mass and foraging guild to infer the framework’s sensitivity dimension (Fig. 1C), and used generation length and clutch size to quantify the adaptive capacity dimension (Fig. 1D). All traits were extracted from a recently compiled dataset of seabird traits (Richards et al. 2021).

### Assessing exposure to bycatch

To estimate the framework’s exposure dimension, we quantified (1) overlap with fisheries activities as the percentage of 1° global grid cells shared between species’ ranges and each gear-specific fishing activity, and (2) fishing intensity as the sum of all fishing hours in the overlapping grid cells (Fig. 1B). To achieve this, we first extracted distribution polygons for 341 seabirds (BirdLife International, 2017) which represent the coarse distributions that species likely occupy, and are presently the best available data for the seabird global ranges. We created a 1° resolution global presence-absence matrix based on the seabird distribution polygons using the package ‘letsR’ and function lets.presab (Vilela and Villalobos 2015). Second, we downloaded the daily fishing effort data for longlines, trawls, and purse seines from Global Fishing Watch, which classifies vessel activity based on vessel type and movements (Kroodsma et al. 2018). For each gear type, fishing effort was summed per 1° global grid cell between 2015 and 2018. Finally, to ensure consistency between the species’ distribution and gear-specific fishing activity layers, we re-projected all spatial data to a raster format with the same coordinate reference system (WGS84), resolution (1° x 1° global grid cells) and extent (± 180°, ± 90°). To achieve this, we used the package ‘raster’ and function rasterize (Hijmans 2020).

### Trait Scoring and Weighting

Each trait, attribute and dimension were scored between 0 and 1, with 1 indicating the greatest vulnerability to bycatch (Potter et al. 2017). This was achieved through a stepwise process. First, all continuous traits from the vulnerability dimensions (body mass, clutch size, generation length, overlap with fisheries, and fishing intensity) were broken into categories using the Sturges algorithm which bins the traits based on their sample size and distribution of values (Sturges 1926). All trait categories were then scored from high to low with ordinal variables based on increased vulnerability to bycatch (Appendix 1-3). To ensure the prioritisation analysis predictably weights the criteria (Mace et al. 2007), all scores were scaled between zero and one and weighted by the frequency of trait occurrence (Potter et al. 2017).

The following worked example represents the scoring and weighting steps for a trait with four categories:

Trait category 1 (lowest vulnerability) = 0
Trait category 2 = (n_1_ + n_2_)/n_total_
Trait category 3 = (n_1_ + n_2_ + n_3_)/n_total_
Trait category 4 (highest vulnerability) = (n_1_ + n_2_ + n_3_ + n_4_)/n_total_ = 1

Where n is the number of species per trait category and n_total_ is the total number of species.

For example, foraging guild contains four categories: ground forager (category 1 = 13 species), generalist forager (category 2 = 63 species), diving forager (category 3 = 121 species) and surface forager (category 4 = 144 species), and n_total_ for this study is 341 species. Ground forager has the lowest conservation priority therefore is given a score of 0. All other foraging strategies are weighted proportionally based on the number of species within that category and the lower categories (Potter et al. 2017). Therefore, generalist forager’s score is (13 + 63) / 341= 0.22, diving forager’s score is (13 + 63 + 121)/ 341 = 0.58 and surface foragers, with the greatest conservation priority, have a score of (13 + 63 + 121 + 144)/ 341 =1. These equations are applied to each trait independently, and the number of trait categories varies between 3 to 5 per trait.

### Vulnerability Classes

We categorise species into vulnerability classes (Fig. 1A) based on a dimension score threshold of 55%. This threshold was decided from a sensitivity test by balancing between excluding all vulnerable species because thresholds were too high, and ensuring minimal species changes between threshold levels across all gear types (Fig. A4.1). If all dimensions (exposure, sensitivity, and adaptive capacity) have a score greater or equal to 55%, species are highly vulnerable to bycatch, therefore, were classified into the “high vulnerability” class. If the scores of sensitivity and exposure were greater or equal to 55%, but adaptive capacity was less than 55%, species were considered to have high vulnerability with potential adaptive capacity, and were assigned to the “potential adapters” class. If the scores of adaptive capacity and exposure were greater or equal to 55%, but sensitivity was less than 55%, species were considered to have high vulnerability with potential to persist and were assigned to the “potential persisters” class. Species were classified into the “potential future vulnerability” class if the scores of adaptive capacity and sensitivity were greater or equal to 55%, but exposure was less than 55%. If all dimensions have a score less than 55%, or if only one dimension has a score greater or equal to 55%, species had low overall vulnerability and were assigned to the “low vulnerability” class. This approach was repeated for the three gear types (longline, trawl and purse seine). Thus, all species received vulnerability scores and classes associated with each gear type.

All analyses were performed in R version 4.0.2 (R Core Team 2020).

## Emerging patterns of species’ vulnerability to bycatch

Our preliminary vulnerability framework revealed emerging patterns within the vulnerability dimensions and classes, with species’ vulnerability varying across the three gear types and dimensions (Fig. 2 & 3; Appendix 5). Albatrosses have the highest overall vulnerability followed by frigatebirds, petrels, and shearwaters, while gulls, terns, and cormorants have the lowest overall vulnerability (Fig. 2). All seabird families have relatively high sensitivity (median = 0.70) and little capacity to adapt (median = 0.74) in response to bycatch (Fig. 2). By contrast, exposure is more variable and has emerged as an important vulnerability dimension. While the median exposure across families is low (median = 0.17; Fig. 2), a number of families and individual species have high exposure scores. For example, the Wedge-tailed Shearwater (*Ardenna pacifica*) has a longline exposure score of 0.95, the Northern Fulmar (*Fulmarus glacialis*) has a trawl exposure score of 0.90, and the Black-tailed gull (*Larus crassirostris*) has a purse seine exposure score of 0.97.

**Figure 2.**
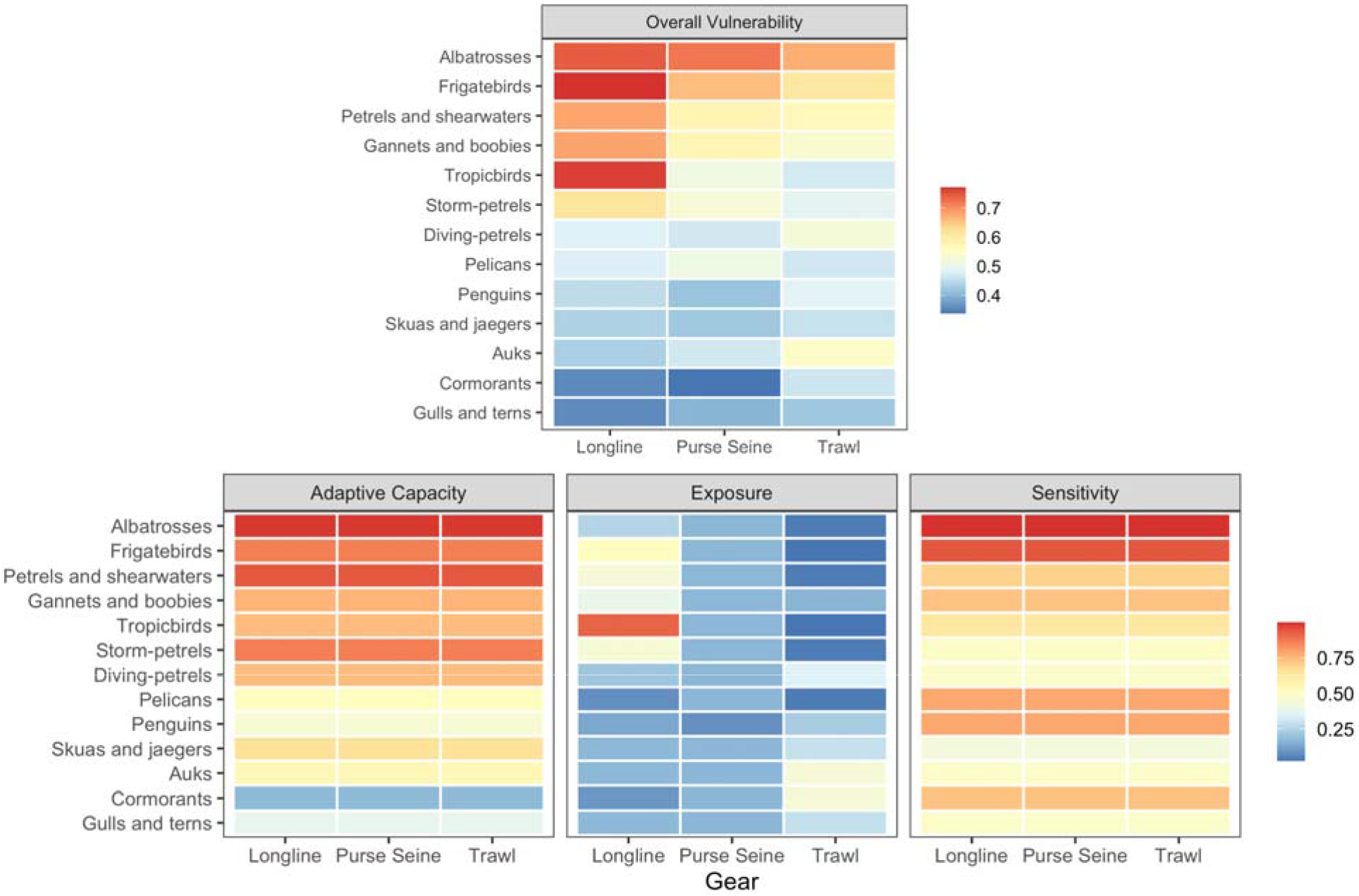
Median overall vulnerability, adaptive capacity, exposure, and sensitivity scores of all seabird families to longline, purse seine, and trawl gear types.

Furthermore, we find 46 species have high exposure (score ≥ 75%) to at least one gear type, but are not identified as vulnerable to bycatch by the IUCN threat classification scheme (threats 5.4.3 & 5.4.4 from https://www.iucnredlist.org/resources/threat-classification-scheme). These species were predominantly gulls and terns (n = 16), petrels and shearwaters (n = 13), and storm-petrels (n = 7). A total of 133 species have lower exposure (score < 75%) to at least one gear type, but are identified as vulnerable to bycatch by the IUCN. These species were predominantly petrels and shearwaters (n = 31), albatrosses (n = 22), auks (n = 19), and gulls and terns (n = 19).

We further find taxonomic differences between the five vulnerability classes. Specifically, species falling into the high vulnerability class (highest scores across all three dimensions) were predominantly albatrosses, petrels, and shearwaters (Fig. 3; Appendix 5). The most frequent species within the potential adapters class (high sensitivity and exposure scores, but do have adaptive capacity due to low scores) were gulls and cormorants (Fig. 3; Appendix 5). Potential persisters (low sensitivity score, high adaptive capacity and exposure scores) were typically storm-petrels and shearwaters (Fig. 3; Appendix 5). The potential future vulnerability class (high scores for sensitivity and adaptive capacity, low score for exposure) was commonly composed of albatrosses, petrels, and shearwaters (Fig. 3; Appendix 5). Finally, species classified with low vulnerability (low scores across all dimensions, or a high score for only one dimension) were predominantly gulls and terns (Fig. 3; Appendix 5).

**Figure 3.**
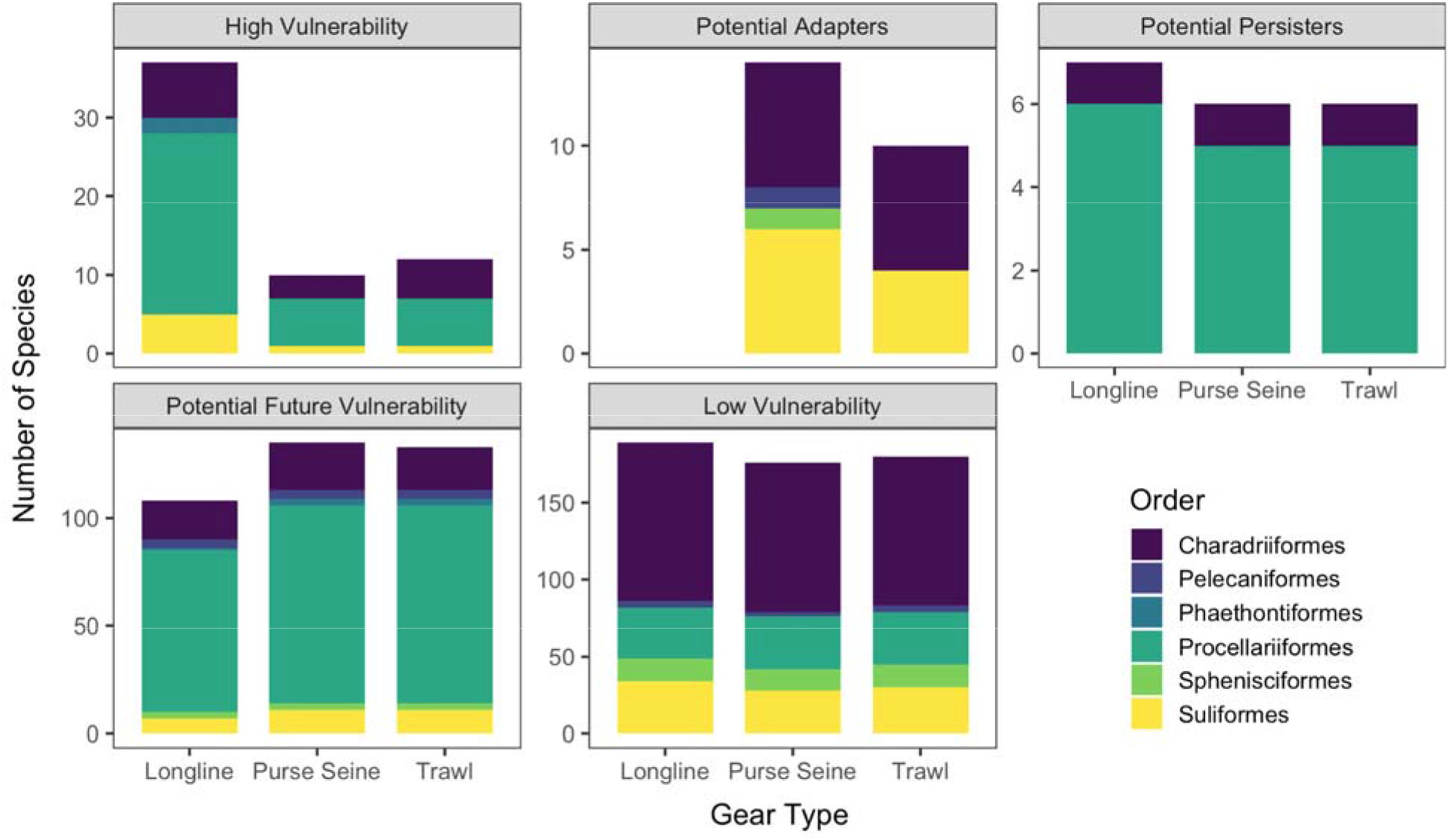
The number of species falling into each vulnerability class for longline, purse seine and trawl gear types. Charadriiforms encompass gulls, tern, skuas, auks, jaegers; Pelecaniformes are pelicans; Phaethontiformes are tropicbirds; Procellariiformes encompass albatross, petrels, shearwaters; Sphenisciformes are penguins; Suliformes encompass gannets, boobies, cormorants, frigatebirds.

## Vulnerability framework limitations

The vulnerability framework identified 62% (n = 32) more species that may be vulnerable to bycatch (those falling into the high vulnerability class), but are not currently recognised by the IUCN threat classification scheme as threatened from bycatch. Furthermore, it is important to note that in its present form, the framework miss-classified 36% (n = 70) of the species identified as threatened from bycatch by the IUCN into the low vulnerability class and 44% (n = 64) into the potential future vulnerability class. These differences are likely attributed to limitations in trait selection within the vulnerability framework’s dimensions. For example, we do not include a species’ boldness or propensity to interact with vessels because these traits are not available for all seabirds. To increase the framework’s value, we encourage its further development in the future with suggestions listed below.

## Future directions for the vulnerability framework

While the framework has been valuable for revealing patterns between and within the vulnerability dimensions, data limitations are presently impeding its full functioning to effectively classify species into their vulnerability classes. However, we believe the framework could become a valuable tool in the future as additional and finer-scale traits and threat data become available because the framework is highly adaptable to spatial and temporal variations in traits and threats. To aid in its replication and development in future analyses, we provide the R code used to build the framework.

### Trait and dimension improvements

While an array of traits are available for seabirds, to strengthen the vulnerability framework’s dimensions, additional efforts are required to compile traits that are not currently available for all seabirds. For example, to improve the sensitivity dimension, future studies may include traits that capture a species’ likelihood of interacting with fishing vessels e.g., boldness, opportunism, competitive ability, and whether they follow ships or not (e.g., Orben et al. 2021). To advance the adaptive capacity dimension, adding additional metrics that relate to breeding and population responses may be important, such as breeding frequency, productivity, and adult survival. Finally, taking advantage of extensive seabird biologging data (e.g. seabirdtracking.org) will be imperative to refine the spatiotemporal resolution of the exposure dimension, through shifting the current fishing overlap metric to a quantification of fishing interaction rate. Moreover, adding information on species abundance distributions and clustering behaviour may further improve the exposure dimension.

### Fishing activity data improvements

Fishing activity and seabird distributions vary daily, seasonally and annually. We therefore acknowledge the limitation of using four years of aggregated fishing activity data. Future modifications of the vulnerability framework may consider integrating the dynamic changes in fishing activity. Moreover, including more gear types could further refine the approach. For example, gillnets fisheries cause an estimated 400,000 seabird mortalities annually (Žydelis et al. 2013). However, we excluded this gear type from our analyses because it presently has poor coverage within the Global Fishing Watch dataset. Finally, distributions of small-scale subsistence, and illegal, unreported, and unregulated (IUU) fishing activities were unavailable, and therefore not included in our vulnerability framework. Incorporating IUU fishing activities in future studies could reveal species with unidentified vulnerability to bycatch.

## A future lens for conservation

Few management actions have incorporated trait-based analyses into conservation strategies (Miatta et al. 2021). However, we suggest that coupling species’ traits with fisheries exposure data within a vulnerability framework could offer an additional lens to advance ongoing conservation measures and policy, such as the IUCN Red List. For example, there is very low observer coverage aboard fishing vessels, and existing data has poor species discrimination and only coarse quantification (Bartle 1991, Weimerskirch et al. 2000, Sullivan et al. 2006, Anderson et al. 2011, Hedd et al. 2016, Suazo et al. 2017). Thus, bycatch mortality of high-risk species may be undetected by on board vessels by fishers and observers, and therefore unreported to the IUCN. The framework could complement vessel-based observations through identifying vulnerable species for which little is known e.g., revealing high vulnerability of gadfly petrels (*Pterodroma* sp.) to longline fleets.

### Local management

This framework could further be extended to inform local management actions. For example, the framework can be easily updated based on interannual and seasonal variation in fishing activity, additional gear types, and reapplied at local scales. We therefore highly recommend future studies couple extensive seabird tracking data with colony-specific trait information and regional fisheries patterns to provide a powerful and informative tool for local management.

## Conclusions

We combined fine-scale fisheries data with seabird traits and distribution data to build a preliminary vulnerability framework that has the potential to identify species at risk from bycatch and help set conservation priorities. Overall, we find most species have high vulnerability scores for the sensitivity and adaptive capacity dimensions. Yet, the framework revealed that species’ exposure to fisheries was highly variable, suggesting that vulnerability to bycatch may be dynamic and rapidly change with future developments in fishing. The framework is highly flexible to trait changes within each vulnerability dimensions, therefore we recommend that future studies compile the additional traits that are required before the framework can be used as a tool to classify species into the five vulnerability classes. Thus, coupling species’ traits with fisheries exposure data within a vulnerability framework could be used as an additional lens to aid ongoing conservation measures and policy. For example, through supporting the efforts of the IUCN Red List and threat identification by suggesting which species need to be especially well investigated and protected.

## Data Sharing and Accessibility

Seabird traits were extracted from (Richards et al. 2021), specifically https://doi.org/10.5061/dryad.x69p8czhd. Species distribution polygons are available upon request from http://datazone.birdlife.org/species/requestdis. Fishing effort data for 2015 and 2016 are available for download, and data for 2017 and 2018 are available upon request from https://globalfishingwatch.org/. Please contact Cerren Richards for access to R code.

