## Supplementary material for "Species’ traits and exposure as a future lens for quantifying seabird bycatch vulnerability in global fisheries": Fig. A4.1

**Appendix 4 –** Sensitivity test

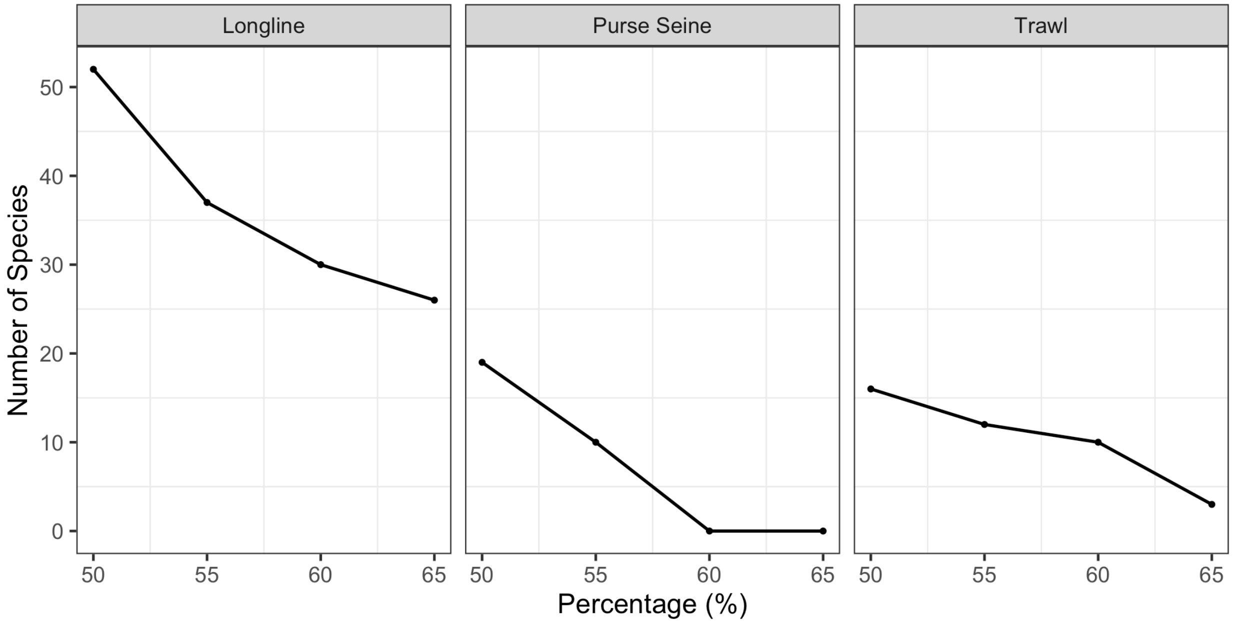


**Figure A4.1** Sensitivity test for the change in number of species per percentage threshold for longline, purse seine, and trawl gear types
